# General mechanisms for a top-down origin of the predator-prey power law

**DOI:** 10.1101/2024.04.04.588057

**Authors:** Onofrio Mazzarisi, Matthieu Barbier, Matteo Smerlak

## Abstract

The ratio of predator-to-prey biomass density is not constant along ecological gradients: denser ecosystems tend to have fewer predators per prey, following a scaling relation known as the “predator-prey power law”. The origin of this surprisingly general pattern, particularly its connection with environmental factors and predator-prey dynamics, is unknown. Here, we explore some ways that a sublinear predator-prey scaling could emerge from density-dependent interactions among predators and between predators and prey (which we call a top-down origin), rather than among prey (bottom-up origin) as proposed in Hatton *et al*. (2015). We combine two complementary theoretical approaches. First, we use phenomenological differential equations to explore the role of environmental parameters and dynamical properties in controlling the predator-prey ratio. Second, we simulate an agent-based model with tunable predator self-regulation to investigate the emergence of predator-prey scaling from plausible microscopic rules. While we cannot rule out alternative explanations, our results show that density-dependent mechanisms relative to predation and intraspecific predator interactions, including prey saturation, predator interference, and predator self-regulation, offer potential explanations for the predator-prey power law.

## 1 Introduction

General biological patterns offer precious opportunities for synthesis and unification [1–8]. Hatton *et al*. [9] put forward evidence of such a general pattern, holding across many terrestrial and aquatic ecosystems, dubbed the “predator-prey power law”. The analysis in Ref. [9] reveals a decrease in the ratio of predator to prey when comparing ecosystems at increasingly higher predator and prey densities. The law holds for focal predator species as well as for aggregate communities and has been recently shown to be robust in ecosystems with more complex trophic structures than previously analyzed (e.g., including omnivores) [10].

In this paper, we try to understand this observation, focusing on a central dataset: the total of large mammalian herbivores and carnivores showing a large gradient of biomass density across a range of 23 African reserves (Fig. 1 and Supplementary Material (SM)). Species composition is similar across the gradient and includes herbivores with a body mass ranging from 5 to 500 kg and large carnivores (20 to 140 kg) [9, 11, 12]. A reason for focusing on this dataset is the robustness of the pattern to various factors, including the taxonomy of species included in the communities, variations in species body mass, the possibility of systematic bias in sampling, and alternative regression approaches [9].

**Figure 1:**
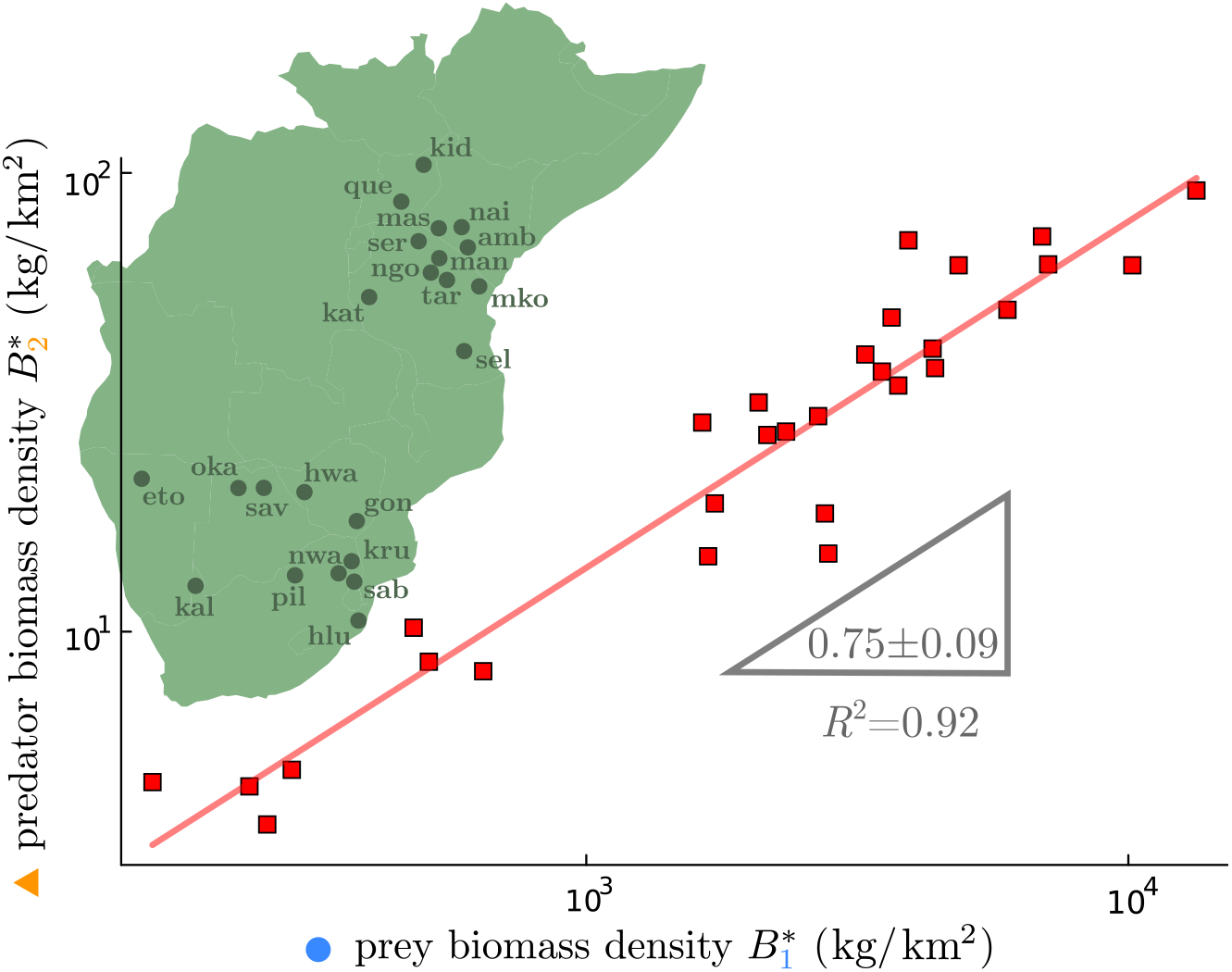
Predator biomass density 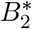 scales sublinearly with prey biomass density 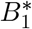, as 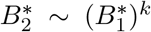 with *k* ≃ 3/4. Data refers to predator-prey communities in African reserves obtained from the meta-analysis in Ref. [9] (see SM). The pattern is robust across ecosystems [9].

Any explanation of this scaling must comprise two parts: what varies between sites to explain that they exhibit different biomass densities and what holds across sites to explain the existence (and exponent) of the scaling law between predator and prey biomass density (see Fig. 2 for a summary of the following argument).

**Figure 2:**
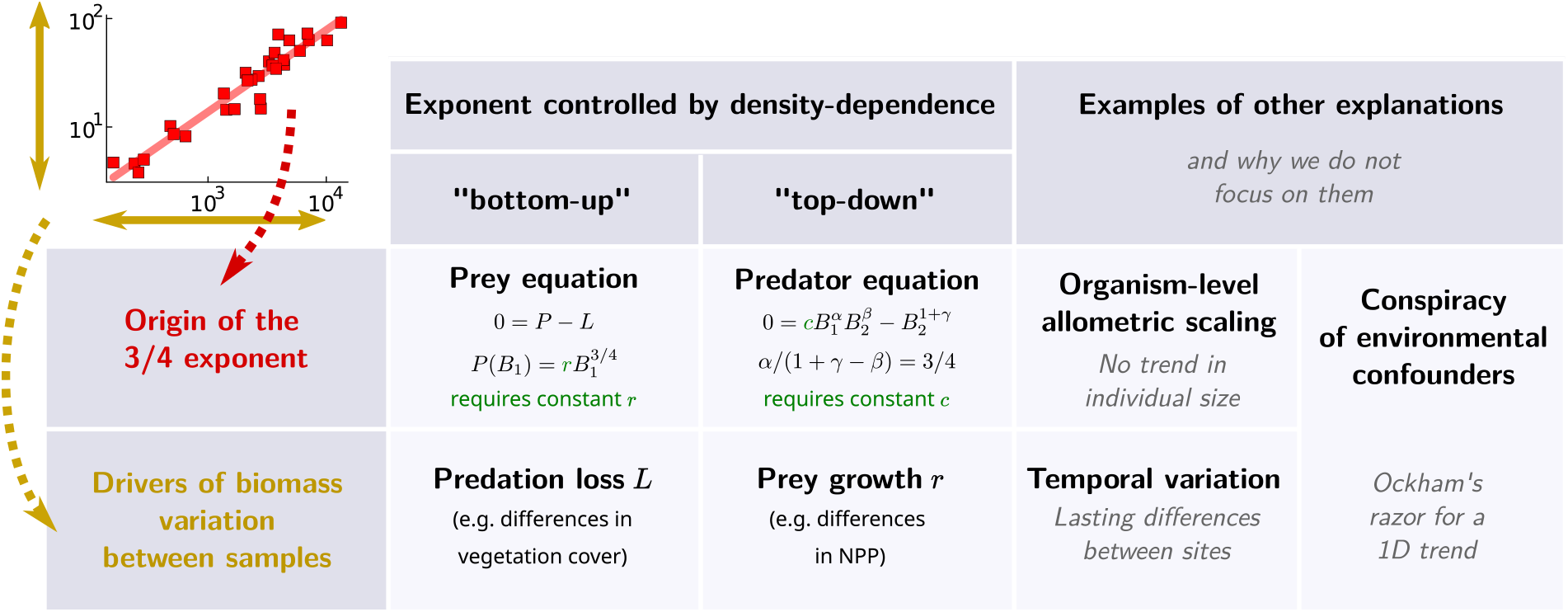
Summary of potential explanations for the predator-prey power law. We need two parts for any explanation: identify the drivers of biomass variation between samples and describe what holds across samples that give origin to the 3/4 scaling. In this work, we focus on explanations in which the 3/4 exponent stems from the functional form (density-dependence) of population dynamics, and choose to call them “bottom-up” when prey consumption and growth control the exponent, and “top-down” when predator consumption and mortality control it. Typically, the source of variation between sites must then lie in the other equation, meaning that the variation in biomasses is expected to arise from ‘bottom-up’ factors, i.e. prey growth, if the exponent of the scaling law is driven by ‘top-down’ factors, i.e. predator consumption, and conversely. We give some examples of other explanations for the predator-prey power law, and of reasons why we are setting them aside here: for instance, the exponent may come from individual-level metabolic scaling, but average organism size and mass do not differ significantly between sites; the differences in biomasses may stem from sites being out of equilibrium, but these differences seem to be lasting (biomass-poor sites do not appear to be simply early stages in biomass growth); the scaling may be due to several parameters changing in correlated ways, but this would require a precise “conspiracy” of physiological and environmental covariates to lead to such a robust, seemingly unidimensional, gradient across sites.

We focus first on what could cause the stark differences in density between different sites. Here, we assume that these densities are close to their equilibrium value given the abiotic and biotic environment (rather than, for instance, low and high densities representing early and late stages in a colonization process), as suggested by the analysis of time series reported in Ref. [9]. Under this assumption, we could start by imagining at least two very distinct explanations.

One possibility is that the gradient is driven by environmental differences among the parks, therefore, by variation in the conditions in which the species find themselves interacting. The other possibility is that, in equivalent environmental conditions, there are differences (phenotypic differences) in the species in the different parks. In the latter case, reaching different equilibria would require having different species or phenotypes, e.g. variation in body mass, size of territory or home range, resource needs, etc. Yet, evidence reported in Ref. [9] stands against the idea that the different parks have significantly different composition in species or individual size, at least none that would systematically create such a gradient. Therefore, we assume that the biomass density gradient is caused by environmental factors.

We can proceed now to explore the reasons behind the specific form of the scaling along the biomass gradient. We first rule out individual-level metabolic theory (which relates organism mass and metabolism or growth with a 3/4 exponent [5, 13]) as the explanation for the community-level 3/4 law, again because there appears to be no systematic variation in individual body mass distribution across the gradient so these metabolic properties should be constant. Turning to other explanations, predator and prey densities could both be controlled entirely by changes in external factors, such as more or less productivity and mortality due to aridity or landscape features, irrespective of any dynamics within and between these trophic levels. In that case, a scaling law between predator and prey density, where both increase in a robust but sublinear relationship, would require a “conspiracy” of parameters (a strong but nonlinear covariation in how the changing environment impacts various species’ productivities and mortalities) for which no ecological mechanism has been proposed yet. Thus we wish to posit that the observed variation of both predator and prey densities stems from the same environmental gradient, but the scaling law between them stems from dynamical mechanisms.

A first explanation of this kind was proposed by Hatton *et al*. [9], suggesting that the driving dynamical mechanism is a sublinear relationship between prey density and prey biomass production. As we recall below, the environmental variation implied by their explanation is that different sites must have different interaction strengths between predator and prey [9], e.g. due to different vegetation cover.

We focus on another ecologically-motivated assumption, proposing that the various parks differ most importantly by their primary productivity, which is positively correlated with herbivore growth (secondary productivity) [14]. Annual precipitation is a good proxy for primary productivity in African parks [15]. In Fig. 3 (**a**), we report predator biomass density vs. precipitation, scaling with an exponent around 5/4 and prey biomass density vs. precipitation, scaling with an exponent close to 3/2 (consistent with Ref. [15]). This results in predator-prey biomass ratio vs. precipitation scaling as −1/4 (Fig. 3 (**b**)). Within the assumption of primary productivity as main driver for the gradient, we will show that a range of explanations are possible, and they are all top-down, in the sense of relying on predation dynamics, and specifically on density-dependent predator interactions. Figure 2 summarizes the explanations for the predator-prey power law we consider in this work and clarify our definition of “top-down” and “bottom-up” mechanisms.

**Figure 3:**
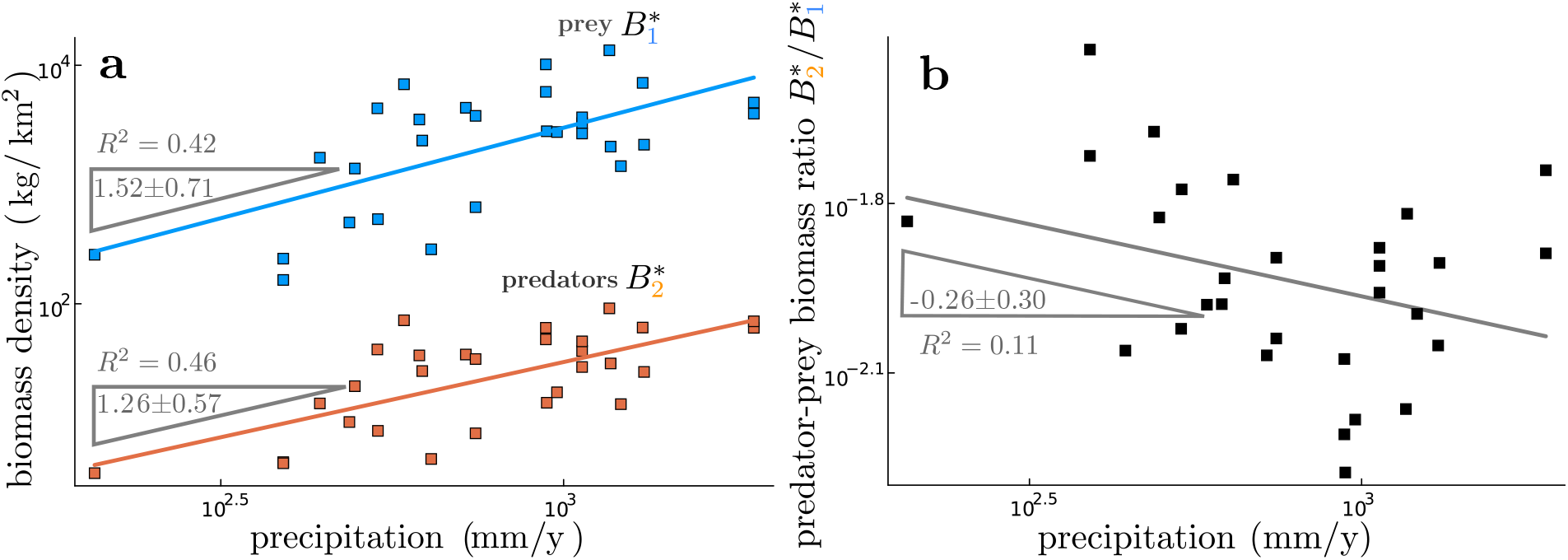
Predator and prey biomass densities scale differently with respect to park precipitation, with their ratio scaling approximatively as −1/4, suggesting that precipitation may be a good proxy for the relevant environmental parameter responsible for the biomass density gradient across different parks. In (**a**), we report predator and prey biomass density vs. park precipitation and, in (**b**), their ratio, again vs. precipitation.

## 2 Predator-prey dynamics and macroecological patterns

We aim to compare ecosystems hosting the same predator and prey species pairs at different stationary biomass densities. We encode the parameters responsible for the biomass gradient in the vector 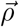. These parameters can represent biotic and abiotic factors affecting the population dynamics (e.g., amount of primary resources or shelters) but also phenotypic traits, which are sensitive to a change of environment but decoupled from the population densities. We collect parameters of the model that are considered independent of the species’ density and do not change across environments in the vector 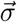. Finally, we define the species density vector 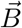 : all the model ingredients representing the dynamics in a given environment are functions of it. We consider a two-dimensional system with one prey species whose density is denoted by *B*_1_, and one predator species, *B*_2_, that feeds on it. This coarse-grained setting allows us to highlight how density dependence clearly can shape the trophic structure at different biomass densities, even though it is not suited to describing a realistic food web in detail.

The equations describing the dynamics of the density of prey *B*_1_ and of predators *B*_2_ can be given, with some degree of generality, by

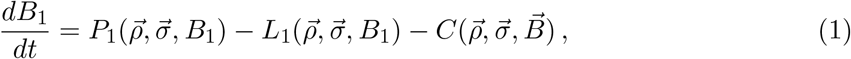

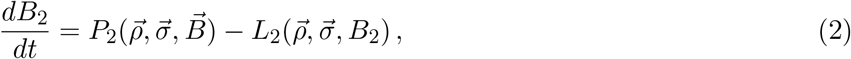

where *P* stands for production, *L* for internal losses and *C* for predation losses. In general, predation losses and predator production depend on both species.

Different choices of what enters in 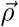 have distinct ecological meanings. A change, e.g., of a coefficient in the interaction term could, for example, represent a different concentration of shelter for prey [16]. Another common reason behind different biomass densities of predators and prey in different environments is a variation in the energy influx in the system through primary production, which can be encoded in a changing coefficient in the prey production term. The observed gradient might not be due to variation in one parameter alone, and we must carefully consider how covariance between the relevant parameters may affect our model predictions [17]. Consider a choice of the production and losses that correspond to the Lotka-Volterra model with self-regulation,

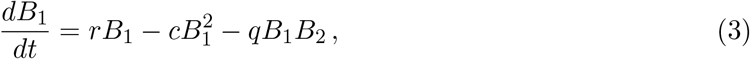

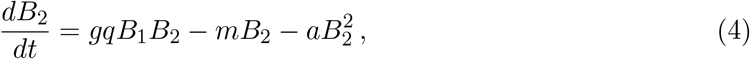

where *r* is the growth rate of the prey, *q* is the interaction strength, *g* the conversion rate, *m* the death rate of the predators, and *c* and *a* the self-regulation strength of prey and predators respectively. It is instructive to write down the stationary solution for the two species’ densities to highlight the explicit dependence on the parameters

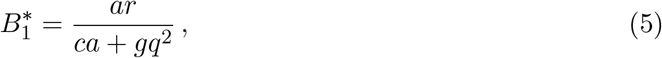

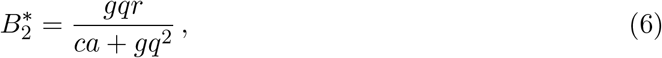

where we considered negligible predator mortality, *m* = 0, for clarity. If the prey growth rate *r* is the single most relevant environmental parameter that changes along the biomass gradient, i.e., *ρ* = *r* and 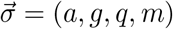, then linear scaling between predator and prey density 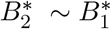 emerges, both being linear in *r*. If, however, the interaction strength *q* is also relevant and it co-varies with *r*, for example as *q* ∼ *r*^*k*−1^, with *k* < 1, then the observed scaling is sublinear 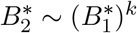.

In summary, the choice of the parameters that are supposed to model a biomass gradient has to be informed by observation, and different choices of parameters produce different macroecological patterns starting from the same dynamical model. We want to explore how dynamical, density-dependent effects can be relevant for the scaling. Therefore, we use as a working assumption that only one parameter, possibly different in other datasets, is responsible for the gradient.

Within this working assumption, a linear relation and, therefore, a constant average number of prey per predator in a patch across a biomass gradient would imply that only the stationary number of predators and prey matters, and how densely packed the two species are is irrelevant. If species density matters [18–21], a sublinear scaling is expected when the net effect of crowding penalizes more the standing biomass of predators than that of prey. In Fig. 4 we pictorially summarize this observation and show examples of mechanisms that could dynamically lead to sublinear scaling. The bottom-up explanation, in the sense of relying on prey dynamics and specifically on density dependence in prey production, emerges from sublinear growth of herbivores [9]. The main classes of mechanisms behind the top-down explanation are predator density-dependent self-regulation, predation interference, and prey saturation.

**Figure 4:**
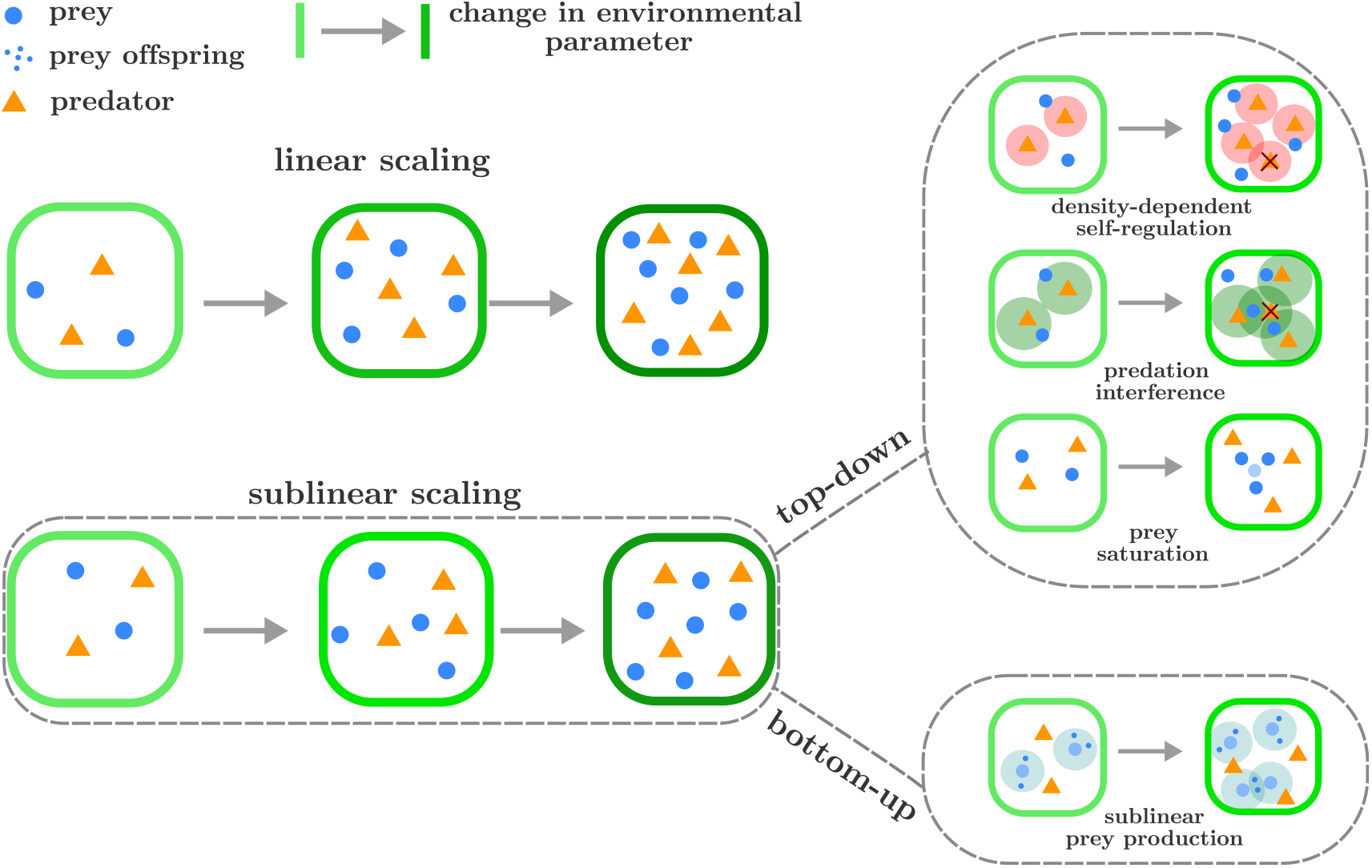
Sublinear scaling reflects the relevance of density-dependent effects for the predatorprey ratio. The figure portrays a *Gedankenexperiment* in which we start with a patch of an ecosystem hosting a given number of predators and prey at stationarity. Then, we change the environmental conditions so that the stationary number of prey is the same in the two cases, but the number of predators scales linearly in one case and sublinearly with the one of prey in the other. Examples of mechanisms that can lead to sublinear scaling are also portrayed, covering bottom-up and top-down explanations. Blue disks represent prey (with small disks representing prey offspring), orange triangles predators. In density-dependent self-regulation (top-down), predators are shown to self-regulate their population size by direct interaction. In predation interference (top-down), on the other hand, this happens because they interfere with each other’s hunting. Prey saturation, shows predators unable to consume all the prey available because some are “shielded” by the others. In sublinear prey production (bottom-up), predators are sustained by prey offspring, and prey are shown to grow sublinearly with their density.

## 3 Bottom-up, sublinear prey production

We summarize here the bottom-up explanation proposed in Ref. [9], which invokes sublinear prey production to describe the observed predator-prey scaling. The model is defined as

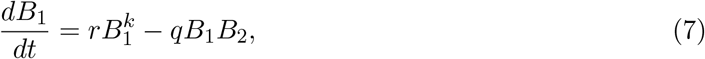

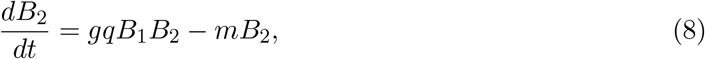

where *r* is the growth rate of the prey, *q* is the interaction strength, *g* the conversion rate, *m* the death rate of the predators, and *k* < 1 is an exponent characterizing the density-dependent effects on the per-capita growth rate of the prey. To be dimensionally consistent, we should multiply the prey production term by a factor 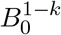, where *B*_0_ has the dimensions of a density. Here, and *mutatis mutandis* in the other sections when not otherwise stated, we consider *B*_0_ = 1 without loss of generality for the purpose of this work.

The equilibrium prey density 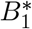 and predator density 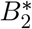, are respectively

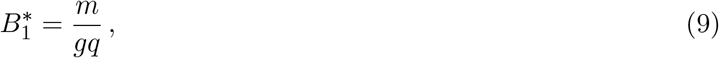

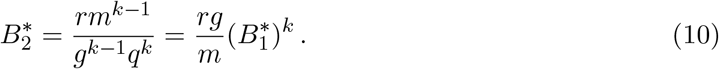

Therefore *rg/m* has to be constant, and *q* has to decrease (increase) to move up (down) across the biomass gradient in order to obtain the scaling 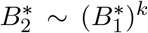. If, instead, *r* is primarily responsible for the biomass increase, this model predicts a constant prey density while the predators grow. In terms of the general definition above, the two cases correspond, respectively, to 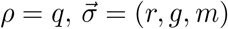 and 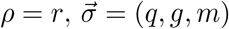.

This bottom-up explanation is informed by another macroecological observation, namely the sublinear scaling of biomass production with respect to biomass density [9, 22]. A discussion on possible mechanistic explanations for sublinear growth can be found in Refs. [9] and [22]; it is, however, still an open question. Sublinear production in the dynamical equation for prey does not represent a unique possible explanation for the macroecological pattern observed across ecosystems. However, it can be realistic and parsimonious, as discussed in depth in Ref. [22].

Support for *q* as the relevant parameter to move along the biomass density comes from cross-system studies, which have shown that density increases across prey causes a decrease of nutritional quality and edibility, which may tend, in turn, to weaken interaction strength with consumers [9, 23–27].

## 4 Top-down, density-dependent predation and predator self-regulation

Here we explore how the predator-prey scaling can have a top-down origin, emerging from density dependence in predation and predator self-regulation. We consider a phenomenological model in which density-dependent effects are important at all scales, following the conceptualization of Ref. [17]. The functional form of the terms involved could, in principle, emerge from underlying spatiotemporal details [28] and should not be necessarily rooted in mechanistic explanations [29]. The model reads

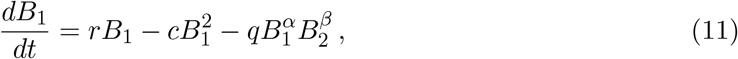

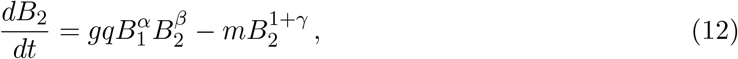

where *r* is the growth rate of the prey, *c* the strength of prey’s self-regulation emerging, e.g., from competition for resources [30, 31], *q, g*, and *m* are, respectively, related to the interaction strength, conversion rate, and predator death rate (by properly accounting for dimensionality, as in the section above). The exponent *α* encodes saturation, *β* is connected to predator interference, and *γ* models the strength of predator self-regulation. The interaction term is scale-free, in the sense that complete saturation with respect to prey density is never achieved. However, this is not unrealistic as evidence suggests that predator feeding rates may often be unsaturated under typical conditions [32]. It is also possible to use other models such as the Hassell-Varley-Holling model [33–36], especially if the systems are far from complete saturation.

In this work, we are only quantitatively interested in the scaling exponents that emerge from the dynamics. Therefore, Eqs.(12) and (11) can be reduced, through non-dimensionalization (see SM), to the form

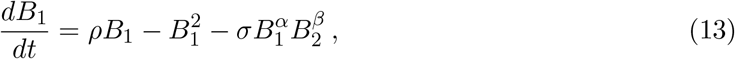

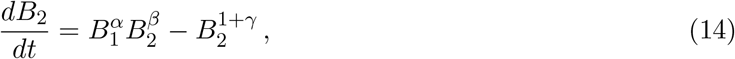

where we call the remaining parameters *ρ* (which is responsible for the biomass gradients) and *σ*. These are functions of the original parameters, with *ρ* ∝ *r* and *σ* independent from *r*. The general stationary solution for predator biomass density as a function of prey reads

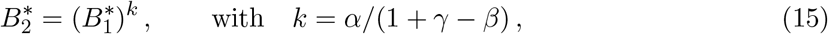

while the solution for the prey has a closed form only for specific values of the exponents (*α, β* and *γ*) and must otherwise be solved numerically. Assuming a varying *ρ*, corresponding to a variation of primary productivity, every combination of the exponents resulting in *k* = 3/4 predator-prey scaling across ecosystems is viable as long as it produces feasible and stable stationary solutions.

As an example, a meta-analysis of predator-prey pairs of mammals, birds, and reptiles suggests that total predation rates increase as the square root of the product of predator and prey density [17], i.e., *α* = *β* = 1/2. In this case, in order to observe the *k* = 3/4 scaling, the exponent associated to predator self-regulation has to be *γ* = 1/6. This means that per-capita losses for predators must scale^1^ like 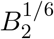. In Fig. 5 we report the predator-prey scaling for this choice of parameters.

**Figure 5:**
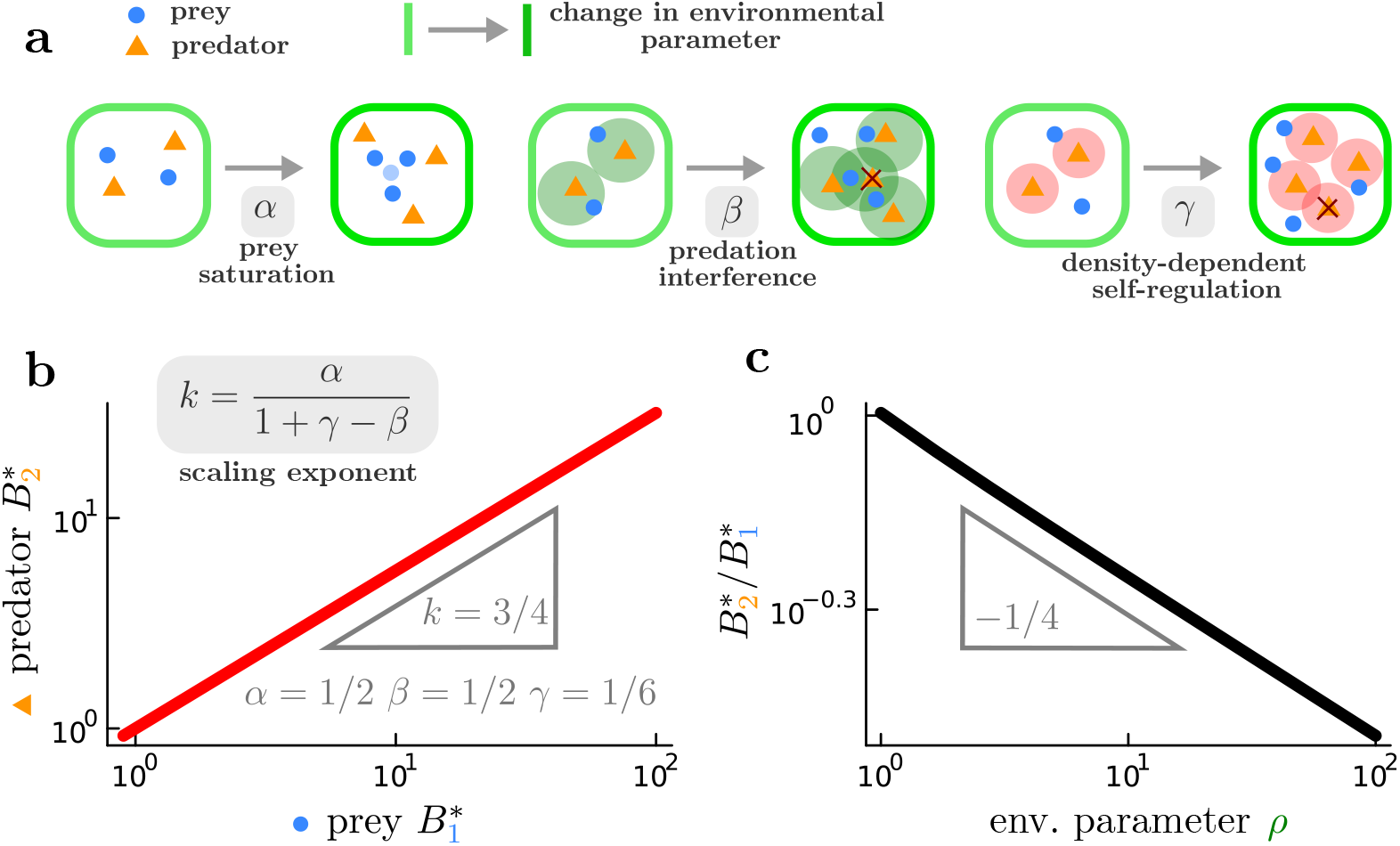
Predator biomass density 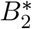 scales sublinearly with prey biomass density 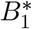 for given exponents for saturation *α*, interference *β* and self-regulation *γ* (**a**) when herbivore growth is varied to simulate a biomass gradient, shown as a straight line on log-log axes. By solving numerically Eqs. (13) and (14), here we show, for *α* = *β* = 1/2, *γ* = 1/6, *σ* = 1/10 and *ρ* ∈ [1, 100], (**b**) the predator stationary biomass density against prey stationary biomass density (*k* = 3/4) and (**c**) their ratio with respect to the environmental parameter *ρ* (see Eq. (13)), which is proportional to the herbivore growth rate *r*.

Differences in primary productivity and herbivore growth are considered among the most common reasons underlying differences in biomass density in otherwise similar ecosystems. Support for considering *r* the relevant variable comes from data on the relation between biomass density and precipitation in the parks (Fig. 3) and from previous studies connecting precipitation with primary productivity and herbivore growth [14, 15]. If precipitation is associated with the relevant variable *ρ* in our model (see Fig. 5 (**b**)), we expect the scaling in Fig. 3. To rule out an obvious potential confounder, we show in the SM that there is no correlation between park area and precipitation level. This evidence is insufficient to establish primary productivity as the most relevant driver, but it indicates it as plausible.

As an additional test, we parameterize this model with realistic values for the focal species in the African parks considered in this work. The scaling relation at stationarity considering the full dimensional model is (see SM)

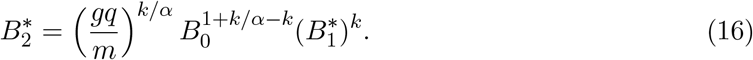

Starting from independent estimations of the parameters *g* [9], *q* [38], *m* [9] and the biomass threshold for nonlinearity *B*_0_ [22], we can estimate the coefficient predicted by our model and confront it with the fitted coefficient from the data *C*_fit_ = 0.075, with dimensions [*C*_fit_] = (kg/km^2^)^1−*k*^ and 95% confidence interval (C. I.) *C*_fit_ = 0.038, 0.15. If we assume *α* = 1, we get

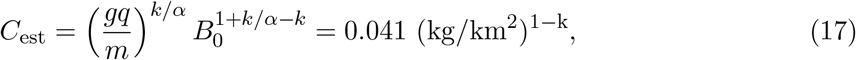

which is within the confidence interval of *C*_fit_. In general *k/α* < 1 is favorable to improve the fit unless *B*_0_ is large. As there is a significant margin for error in all these estimates, this calculation is only meant as a “sanity check” to verify that our model predictions are at least compatible with some empirical evidence.

## 5 Agent-based model as a proof of concept for the top-down case

Predator-prey agent-based (ABM) models [39–43] can provide a bridge between mathematical models and more realistic behavioral rules, providing more intuition on the dynamics underlying our patterns. We present a simple ABM that produces sublinear predator-prey scaling across a biomass gradient through top-down effects. We focus, as an example, on density dependence in predator self-regulation.

We consider a *L* × *L* grid on which the agents move with Moore connectivity (each site has 8 neighbors) and periodic boundary conditions. No occupancy restriction is enforced: there is no upper bound to the number of agents a grid point can host. There are two species: prey and predators. Agents of both species possess an internal energy that changes during the dynamics; metabolism decreases it, feeding increases it, and during a reproduction event, the parent donates half of its energy to the offspring. Prey move on an adjacent grid point, and their energy changes of an amount ∆*E* = −*μ*, where *μ* is the metabolic rate. Then, they eat a fraction of the resources present on the grid, *R*, which is converted into energy ∆*E* = *R/N*_1_ where *N*_1_ is the numbrer of prey present in the grid point. Finally, if their energy is *E* ≤ 0 they die, otherwise, with probability *P* = *ω*_1_ they reproduce. Predators dynamics is slighlty more involved. They move and their energy changes as ∆*E* = −*μ* due to metabolism. Then, with probability *P* = *N*_1_*/*(*N*_1_ + *N*_2_) they engage a prey while with with probability 1 − *P* engage a predator, where *N*_2_ is the number of predators in the grid point. In the first scenario, with probability *P* = *δ*, they eat the prey, gaining ∆*E* = *η*, where *η* is the predation efficiency, and subsequently, with probability *P* = *ω*_2_, they reproduce. In the case in which they do not catch the prey, and if their energy is zero or less, they die. In the scenation in which they engeage a predator, they kill the competitor with probability *P* = *f*. Finally, if their energy is *E* ≤ 0 they die of starvation. At the end of the time step the amount of resources at each grid point increases by *ρ*, the resource growth rate. The functioning of the model is summarized in Fig. 6.

**Figure 6:**
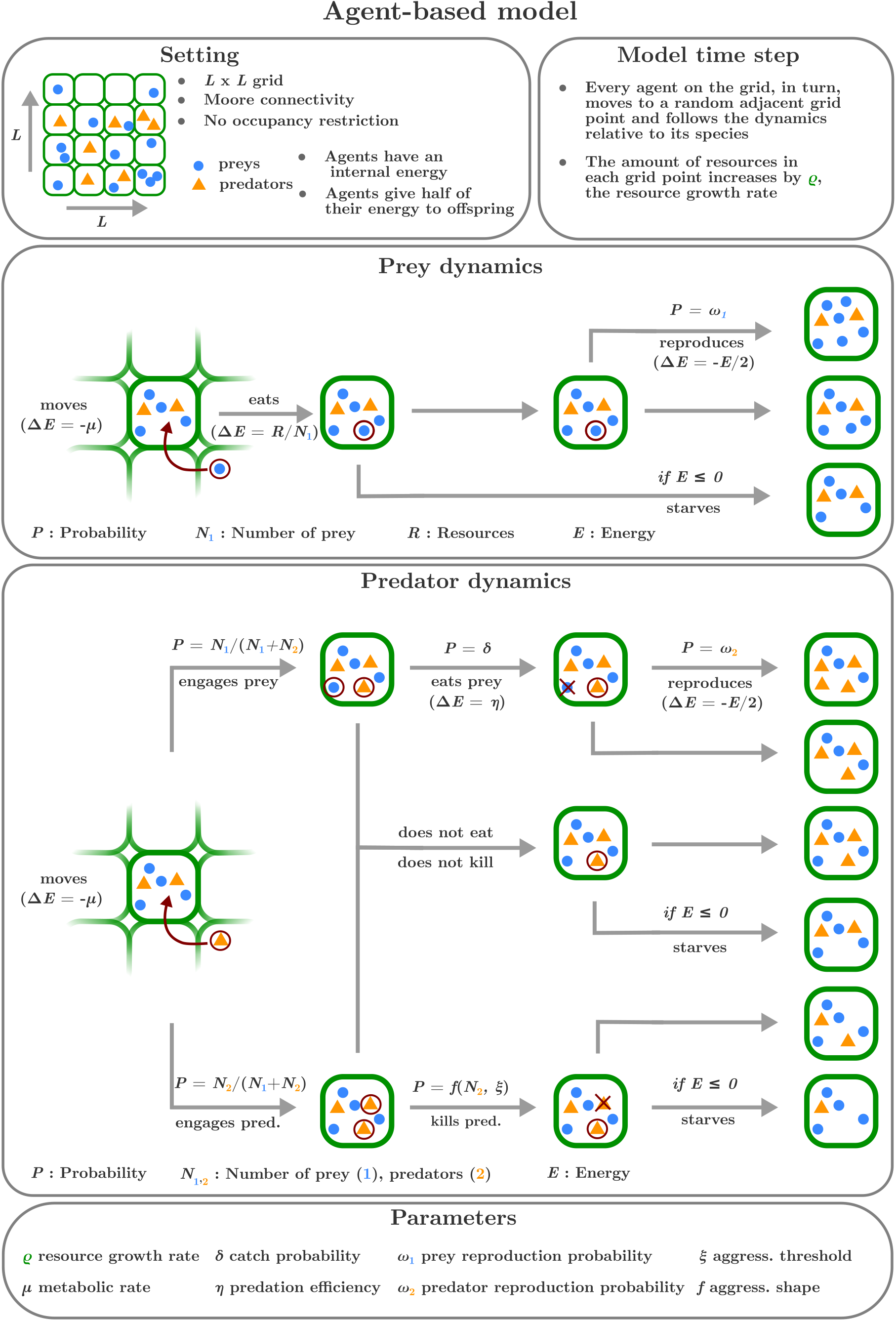
Visual description of the agent-based model. The general setting and the parameters are defined, as well as predator and prey dynamics.

A biomass gradient in the model is obtained by tuning the resource growth rate *ρ*: the higher *ρ*, the higher the biomass density at stationarity. The key element of this model is the density dependence in the probability that a predator has of attacking and killing another predator when encountered. We call it *aggressiveness* and denote it by *f*(*N*_2_, *ξ*), where *ξ* is a parameter that establishes a threshold over which the attack is certain. It controls the sensitivity of this mortality with respect to the predator density (i.e., number at a given grid point). In order to explore more regimes, we study the form

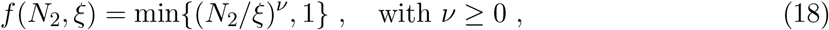

where the exponent *ν* modulates the dependence of the aggressiveness on predator density. We choose for our simulations a threshold *ξ* = 100 that allows for a large enough range of prey densities.

Figure 7 shows the equilibrium biomass density of predators 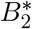 as a function of the equilibrium biomass density of prey 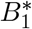 (equivalent to the numerical density upon choosing unitary mass for both species without loss of generality) for different values of the resource growth rate *ρ*. If the aggressiveness does not depend on the number of predators in the grid point (*ν* = 0) a linear scaling is observed. Any positive value of *ν* implies sublinear scaling. The general asymptotic expression of the density of predators as a function of the density of prey reads (see SM)

**Figure 7:**
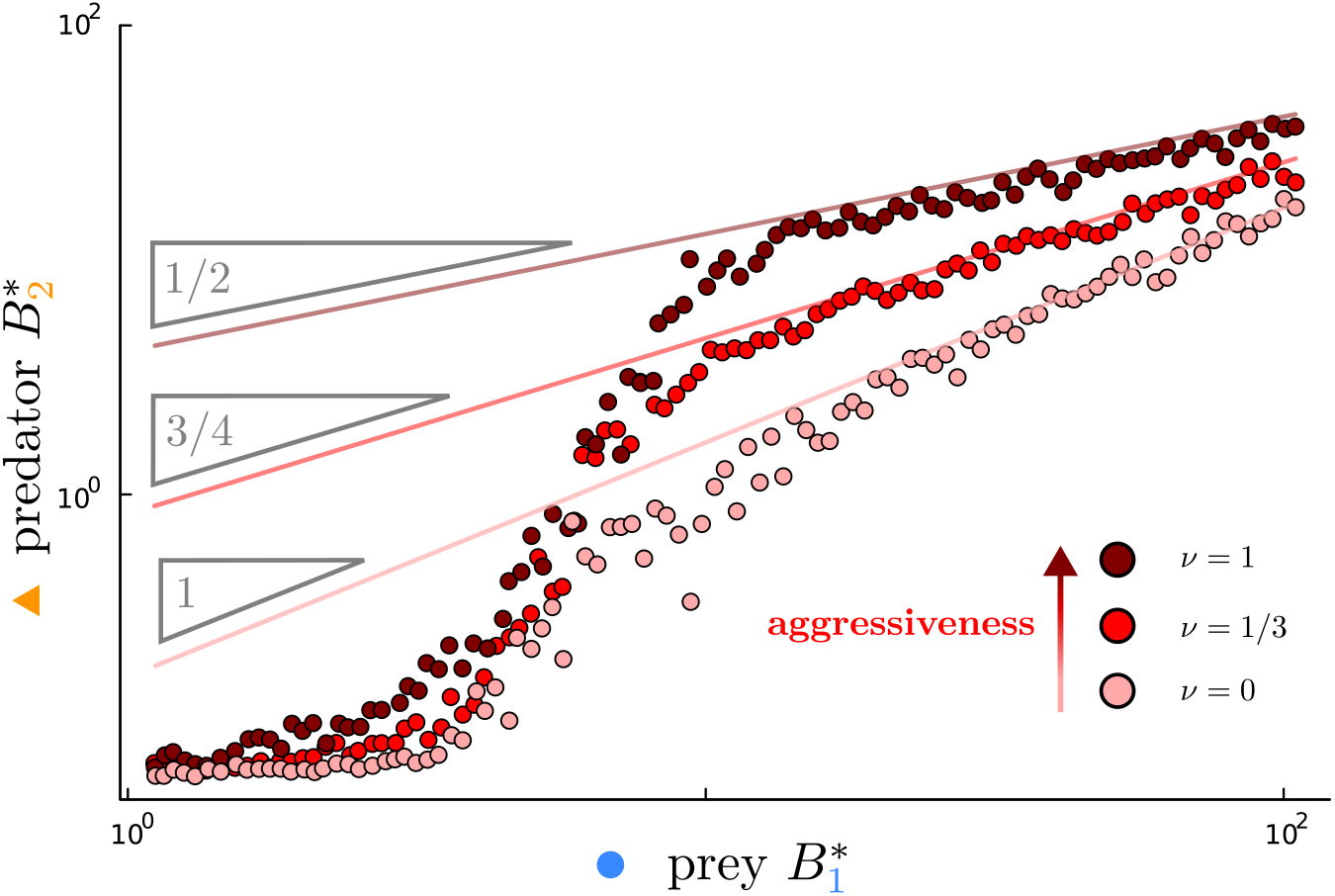
In the ABM, predator biomass density 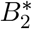 scales more and more sublinearly with respect to prey biomass density 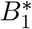 for increasing levels of aggressiveness (encoded in the exponent *ν* and described by Eq. (18)). Each point is a time average of the densities over simulations that reached stationarity (in particular from step 50 to step 500), for each value of the aggressiveness exponent *ν* the resource growth rate *ρ* varies between 1 and 100. The grid area is 4^2^, the initial density of prey and predators is 1, the aggressiveness threshold is *ξ* = 100, the reproduction probabilities are *ω*_1_ = *ω*_2_ = 1*/*2, the catch probability is *δ* = 1/3, the predation efficiency is *η* = 10, and the metabolic rate *μ* = 1. The resulting scaling exponents depend only on *ν* and are robust to changes in the other parameters. The gray lines are theoretical predictions from Eq. (19).

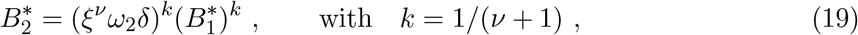

where *δ* is the probability of a catch when a predator engages with a prey.

The model presented here serves as a proof of concept that density-dependent deleterious effects for predators (in this case, self-regulation) can induce sublinear scaling across a biomass gradient. It also shows that the scaling relationship may break even with the right ingredients. In this case, the predator population cannot be sustained below a given resource growth rate, a feature not uncommon in predator-prey models with explicit resource dynamics (see SM).

A comparison with the phenomenological model in Eqs. (13) and (14) is possible through the two expressions for the scaling exponent *k* (Eq. (15) and Eq. (19)) and thanks to the fact that in both models predator density depend on the environmental parameter only through prey density. A parameterization of the phenomenological model that captures the ABM dynamics is *α* = 1, *β* = 1 and *γ* = *ν* + 1.

## 6 Discussion

In this work, we showed that top-down mechanisms represent a viable explanation for the origin of the predator-prey power law, alongside the bottom-up explanation in Ref. [9]. Under the assumption of prey productivity as the primary driver for the biomass density gradient across ecosystems, we demonstrated that combinations of nonlinearities in the interaction term and predator self-regulation can produce the predator-prey power law. While there is no single model that can characterize predator-prey interactions in general [29, 44–46], we considered a scale-free model [17] which capture the relevant dynamical trends that could be behind the macroecological pattern. We also devised and analyzed an agent-based model to provide an example of how predator-prey scaling can emerge as a top-down effect starting from plausible microscopic rules.

We are not able to rule out alternative explanations. Extensive field studies are needed to identify the most relevant factors (and their covariance) behind differences in equilibrium biomass density in ecosystems with similar species composition. This identification is crucial to understand if dynamical ingredients and which ones determine the emergence of predator-prey scaling. We hope to stimulate field work and analysis of existent data sets to test the plausibility of the top-down explanation presented here and, possibly, identify the specific density-dependent mechanisms behind it. Cross-systems studies on predator self-regulation analogous to the one performed for the interaction term in Ref. [17] would be particularly valuable.

It is also possible to put to the test different models by requiring their consistency with other macroecological patterns, such as the sublinear scaling of production with respect to biomass density [9, 22]. The scaling exponent is approximately the same as in the predator-prey scaling and is recovered for prey production in the bottom-up framework. The top-down model predicts, by the equilibrium requirement on the prey (11), that prey production scales as 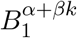 for a range of densities and then as 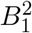 for higher densities, depending on the value of *σ*. For *α* = *β* = 1/2 and *γ* = 1/6 the exponent is 0.875, a bit higher than 3/4. If *γ* = 0, we have the same exponent *k* = *α/*(1 − *β*) for both patterns until production scales as 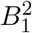 for high enough densities. From this perspective, the bottom-up model is more appropriate to capture the general dynamics. Production scaling with biomass density, however, is not recovered for predators; and although available data are still fragmentary, they are indicative that the pattern should hold at every trophic level [9, 22].

Establishing if either of these two classes of dynamical explanations for the predator-prey power law is close to a correct description of reality, at least at large scales, may also impact how we can act upon these ecosystems. For example, an increase in prey productivity *r* would have a completely different effect on species densities in the two cases. Policies of wildlife management based on the two different dynamical models would result in different ecological responses.

Finally, future works might benefit from the inclusion of evolution. Macroecological scaling patterns in plankton communities can emerge from body size scaling through eco-evolutionary dynamics [47]. This, however, requires producer (phytoplankton) mean body size variation with density, which is not observed in other systems, such as the mammalian herbivores and carnivores analysed in this work.

Data are available in the SM and at [9]. The Julia code used for analysis and simulations is available at https://github.com/onofriomazzarisi/top-down-vs-bottom-up.

## Acknowledgments

We thank J. Arnoldi, A. Bideault, L. Fant, B. Girardot, I. Hatton and M. Novak for fruitful discussions and critical feedback on the work. Funding for this work was provided by the Alexander von Humboldt Foundation in the framework of the Sofja Kovalevskaja Award endowed by the German Federal Ministry of Education and Research, by the Trieste Laboratory on Quantitative Sustainability - TLQS and by NSF BIO OCE grant 2023473. A preprint version of this article has been peer-reviewed and recommended by PCIEcology (https://doi.org/10.24072/pci.ecology.100692).

## Supplementary Material

### 6.1 Data on African communities

In this section, we report the details and references for the data reported in the paper relative to African communities obtained from the meta-analysis provided in Ref. [9]. Moreover, we present further analysis of the same data.

The data chosen as representative of predator-prey systems used to produce the Figures in the main text are reported in Tab. 1. Table 2 associates the full park names to the park codes reported in Tab. 1. Figure 8 reports the behavior of predator vs. prey biomass density, biomass, number density, and number, Fig. 9 reports predator and prey biomass and number density vs. park area, Fig. 10 reports predator and prey biomass and number density vs. precipitation and Fig. 11 reports precipitation vs. park area.

**Table 1:**
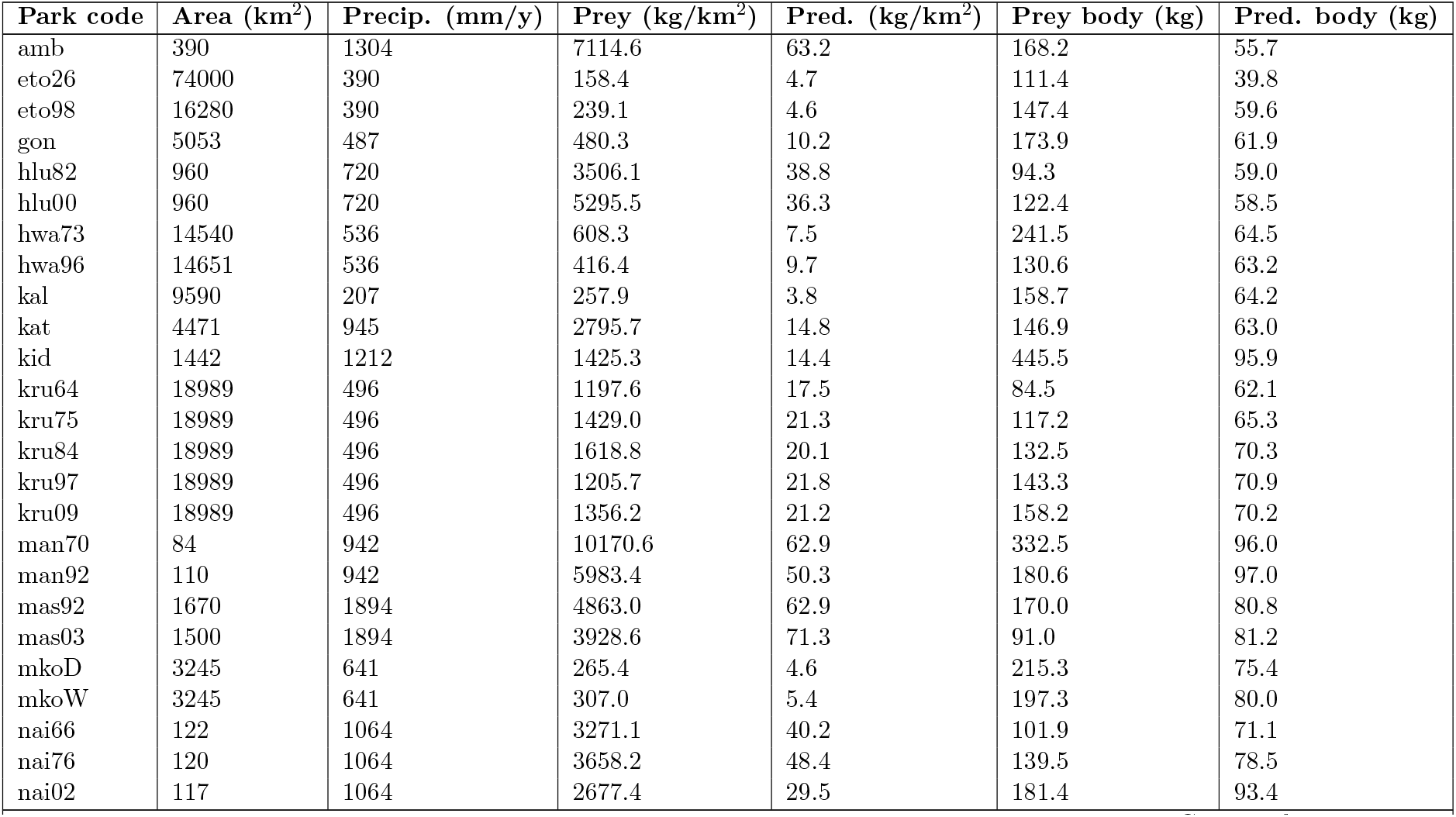
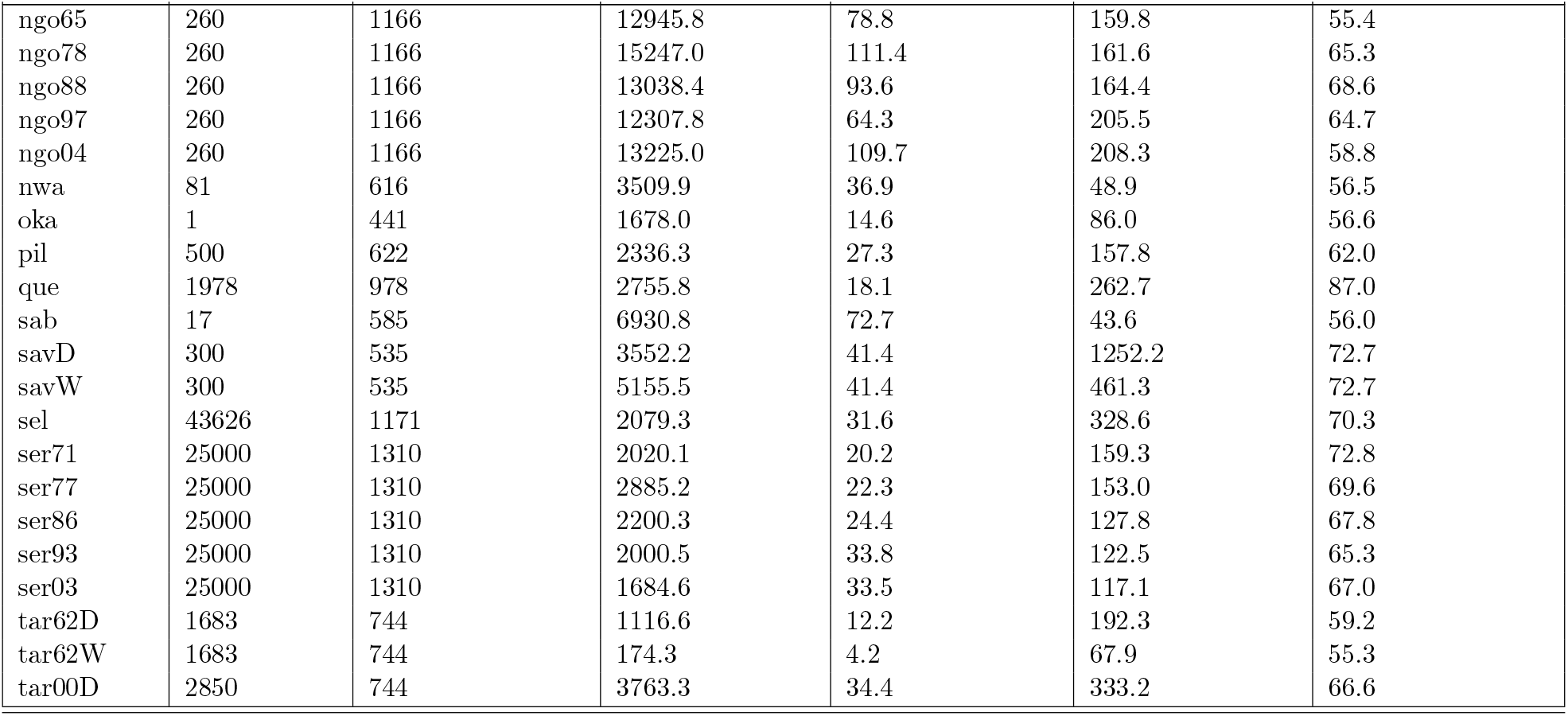
Data for African communities obtained from Ref. [9], In particular, these are the aggregated data for predator species and prey species for African national parks, for details and further references see Ref. [9]. For the plots in the main text, the data referring to the same park (when sampled with the same size, e.g., this does not apply for eto26 and eto98) are averaged. The number in the code refers to the year of the measurements, and the D and W refer, respectively, to measurements in the dry and wet seasons. Correspondence between the park code and the park’s full name is reported in Table 2.

**Table 2:** Park corresponding to park codes.

| **Park code** | **Park** |
| --- | --- |
| amb | Amboseli National Park, Kenya |
| eto | Etosha National Park, Namibia |
| gon | Gonarezhou National Park, Zimbabwe |
| hlu | Hluhluwe iMfolozi National Park, South Africa |
| hwa | Hwange National Park, Zimbabwe |
| kal | Kalahari National Park, South Africa |
| kat | Katavi National Park, Tanzania |
| kid | Kidepo Valley National Park, Uganda |
| kru | Kruger National Park, South Africa |
| man | Lake Manyara National Park, Tanzania |
| mas | Masai Mara National Reserve, Kenya |
| mko | Mkomazi Game Reserve, Tanzania |
| nai | Nairobi National Park, Kenya |
| ngo | Ngorongoro Crater, Tanzania |
| nwa | Nwaswitshaka River, Kruger NP, South Africa |
| oka | Okavango Delta, Botswana |
| pil | Pilanesburg National Park, South Africa |
| que | Queen Elizabeth National Park, Uganda |
| sab | Sabie River, Kruger NP, South Africa |
| sav | Savuti area of Chobe National Park, Botswana |
| sel | Selous Game Reserve, Tanzania |
| ser | Serengeti ecosystem, Tanzania |
| tar | Tarangire National Park, Tanzania |

**Figure 8:**
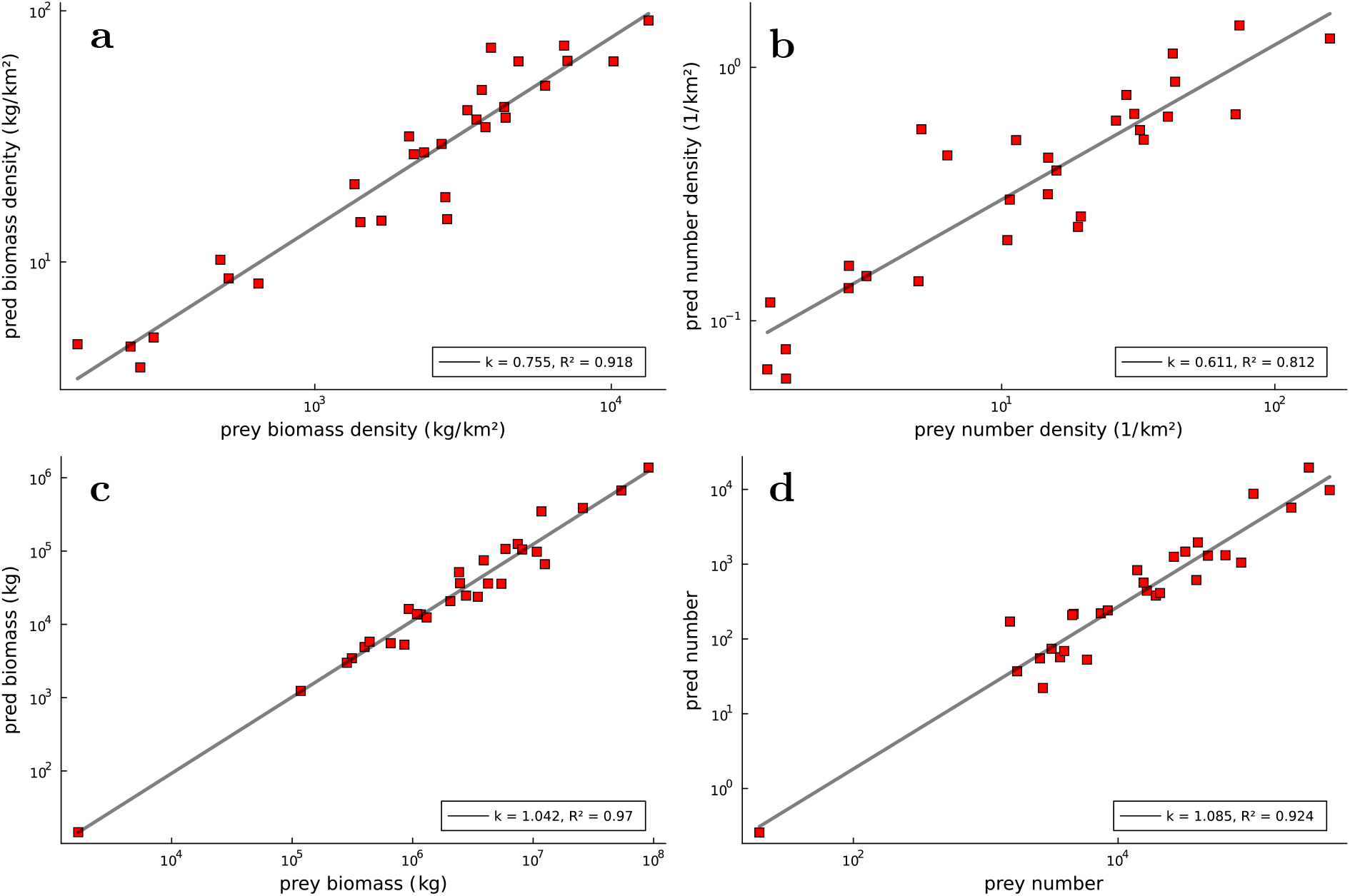
Predator vs prey biomass density (**a**), number density (**b**), biomass (**c**) and number (**d**).

**Figure 9:**
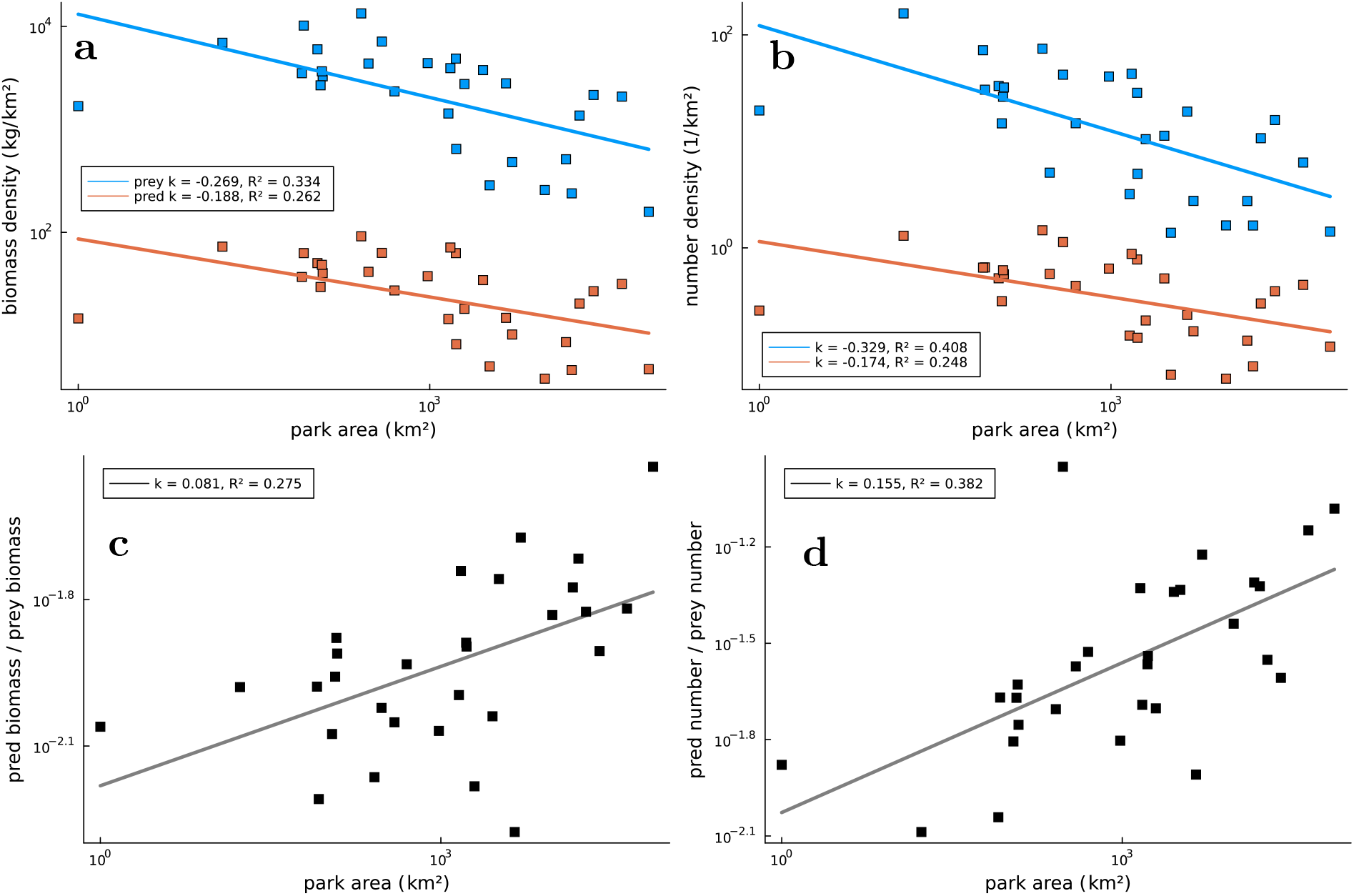
Predator and prey biomass density (**a**) and number density (**b**) plotted against park area. Their ratio is plotted again vs. park area in (**c**) and (**d**).

**Figure 10:**
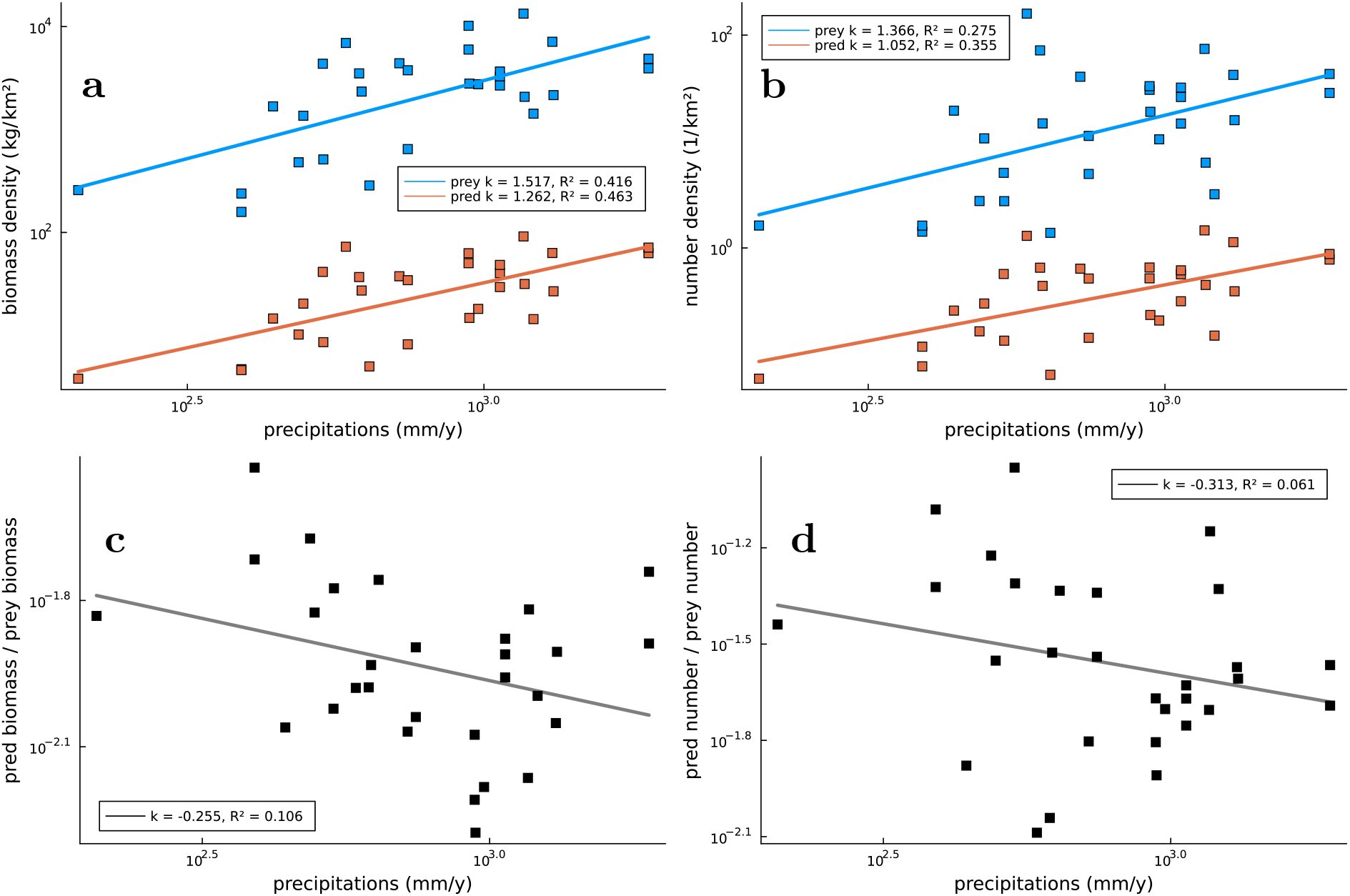
Predator and prey biomass density (**a**) and number density (**b**) vs. precipitation. Their ratio is reported in (**c**) and (**d**).

**Figure 11:**
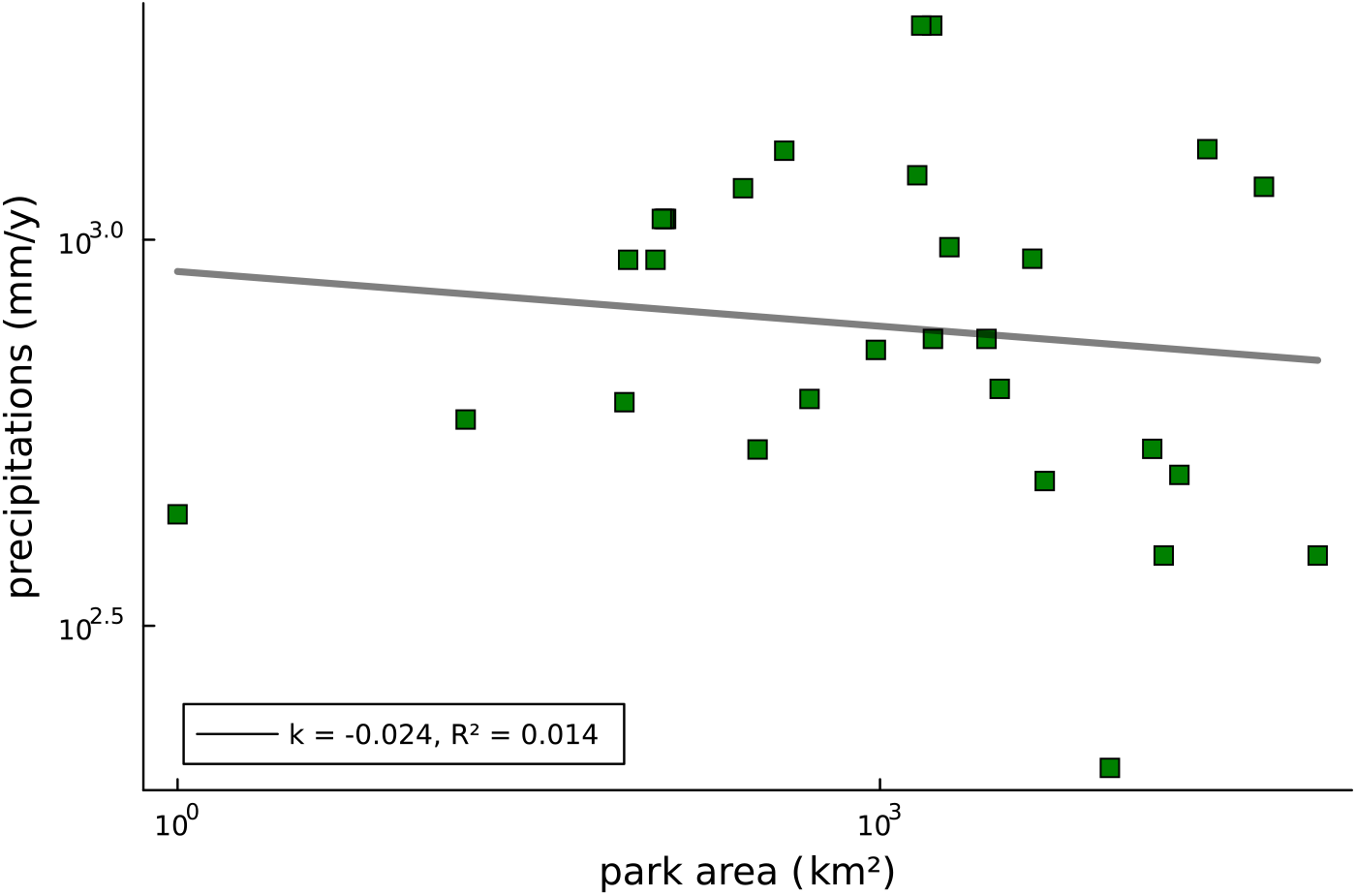
Park precipitation is not correlated with park area.

### 6.2 Phenomenological model analysis

In this section, we provide details of the analysis of the scale-free phenomenological model discussed in the main text to explore density-dependent effects in predation and self-regulation.

We start by rewriting the model in non-dimensional form. The model describes the dynamics of prey biomass density (*B*_1_) and predator biomass density (*B*_2_) through the following equations

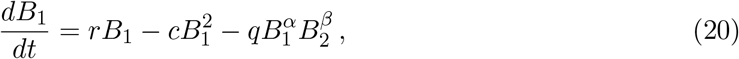

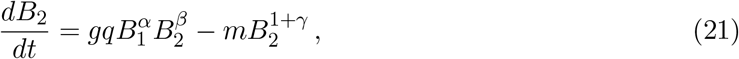

where we refer to the main text for the definition of the parameters. Defining the nondimensional quantities

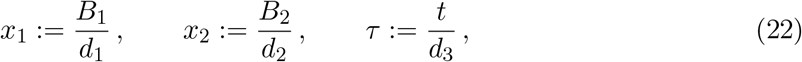

The equation above can be written as

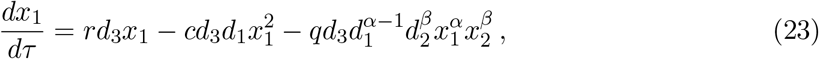

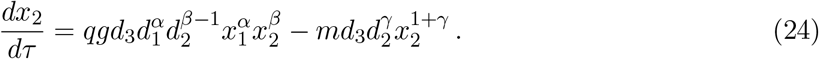

If we make the choices

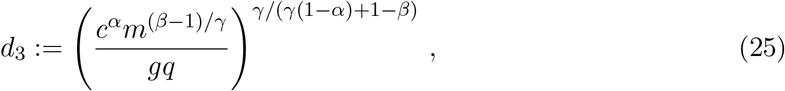

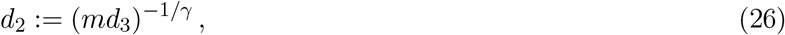

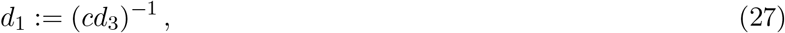

the non-dimensional equations become

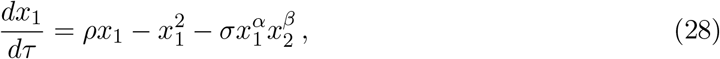

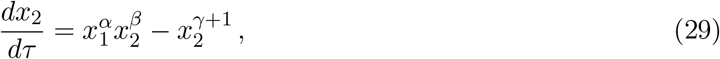

with

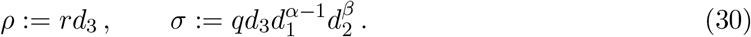

Notice that *r* does not appear in the definition of *d*_1_, *d*_2_ and *d*_3_. Therefore, when varying *r*, and therefore *ρ* as in the main text, we can be sure of the absence of spurious correlations hidden in the redefinition of the densities and time and in the definition of *σ*.

### 6.3 Parameterization of the phenomenological model

In this section we do a sanity check on the phenomenological model with parameters extracted from literature. Let us consider the full phenomenological model, including dimensional constants we ignored in the main text

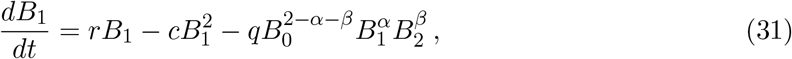

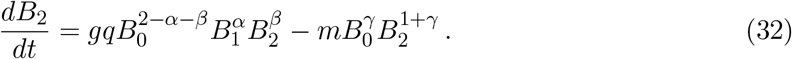

The scaling relation at stationarity is

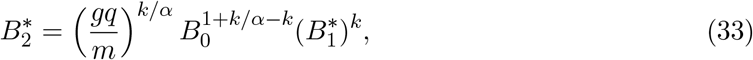

where *k* = *α/*(1 + *γ* − *β*). The fit of the data gives

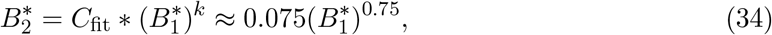

where *C*_fit_ has dimensions [*C*_fit_] = (kg/km^2^)^1−*k*^ and 95% confidence interval (C. I.) *C*_fit_ = 0.038, 0.15. We can use independent estimates of *g, q, m* and *B*_0_ to estimate the coefficient and confront it with *C*_fit_. The dimensionality of the parameters is [*g*] = 1, [*q*] = km^2^/(kg day), [*m*] = 1/day and [*B*_0_] = kg/km^2^. We estimate *g* = 0.0083 [9], *m* = 0.00023 1/day [9] and *B*_0_ = 10 kg/km^2^ [22]. We can decompose the parameter *q* as *q* = *s/b*_2_, where *s* is the search rate and *b*_2_ is the predator body mass. In Ref. [38], the authors find for interaction taking place in 2 spatial dimensions the following scaling relationship

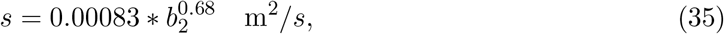

where the predator body mass, *b*_2_, has to be expressed in kg. By using the mean body mass among the predators in all the parks *b*_2_ = 70 kg and expressing *s* in km^2^/day, we get a search rate of *s* = 0.00129 km^2^/day, and therefore we can estimate *q* = 0.000018 km^2^/(kg day). Finally, if we assume *α* = 1, we estimate

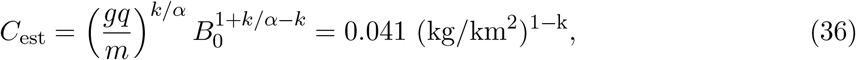

which is within the confidence interval of *C*_fit_ (95% C. I. *C*_fit_ = 0.038, 0.15).

### 6.4 Agent-based model analysis

In this section, we derive the asymptotic expression Eq. (19) for the stationary predator density as a function of the prey density across a biomass gradient.

In our ABM, the probability for a predator to reproduce is given by the probability of engaging and eating a prey times the probability of giving birth. Moreover, at high densities, predators mostly die because of attacks from other predators and not from starvation. We can, therefore, write an equation for the evolution of the density of predator *x*_2_ valid at high densities

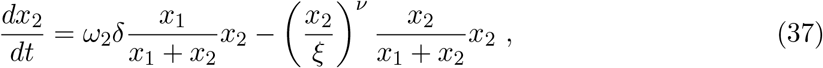

where *x*_1_ is the prey density, *ω*_2_ is the predator reproduction probability, *δ* is the catch probability, *ξ* is the territoriality constant and *ν* is the aggressiveness exponent. At stationarity, we have

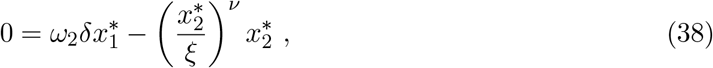

and therefore Eq. (19)

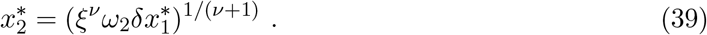

In Figs. 12, 13 and 14 we report the effects of changes in the parameters *η μ* and *ω*_1_ which do not appear in the scaling Eq. (19) on the ABM dynamics at fixed *ρ*.

**Figure 12:**
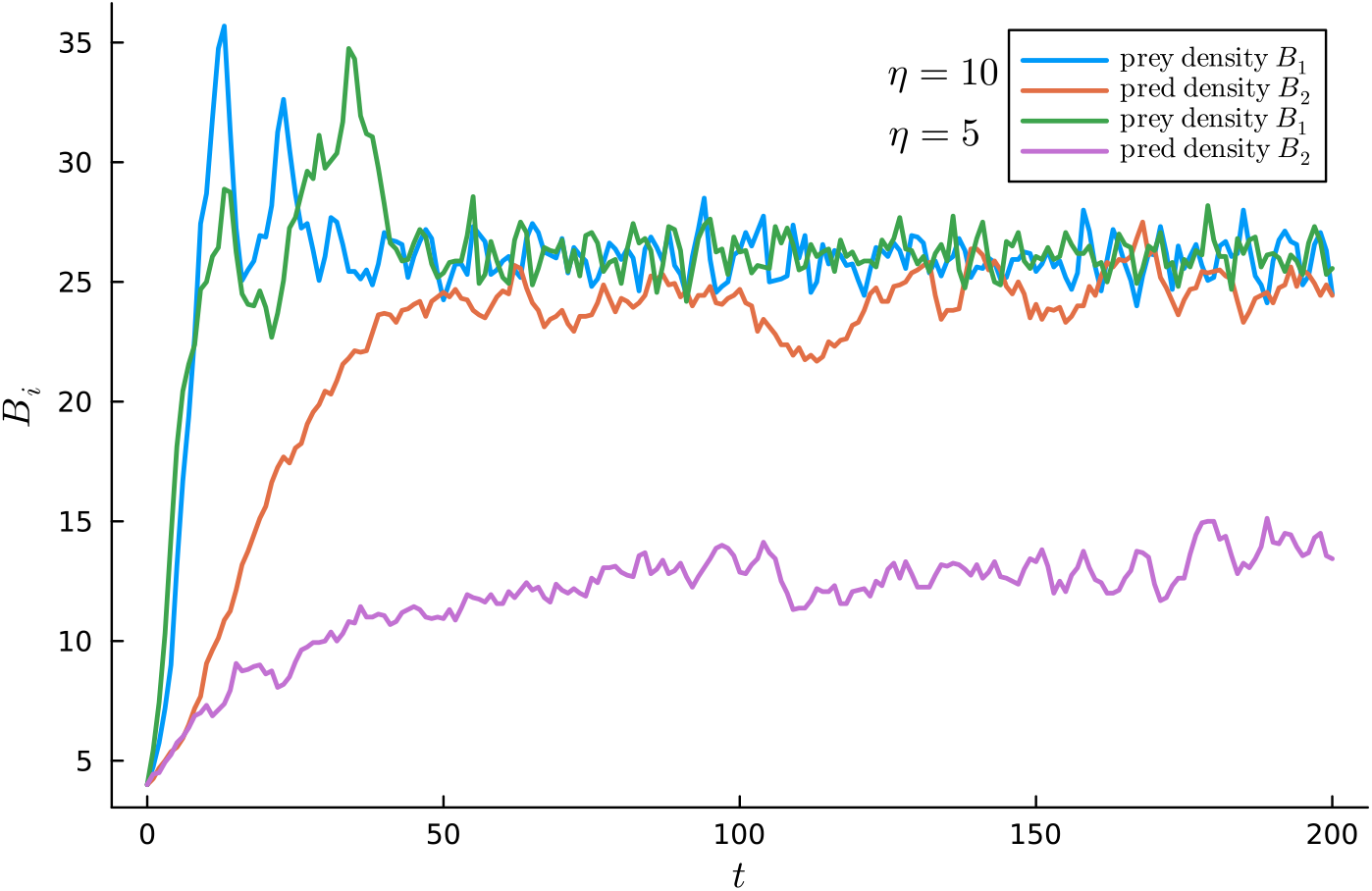
In the ABM, decreasing the value of the predation efficiency *η* decreases predator stationary density and does not affect prey stationary density. The grid area is 4^2^, the initial density is 1 for prey and 10 for predators, the aggressiveness threshold is *ξ* = 500, the reproduction probabilities are *ω*_1_ = *ω*_2_ = 1/2, the catch probability is *δ* = 1/3, the resource growth-rate is *ρ* = 25, the metabolic rate is *μ* = 1 and the predation efficiency is *η* = 10 for blue and orange and *η* = 5 for green and purple.

**Figure 13:**
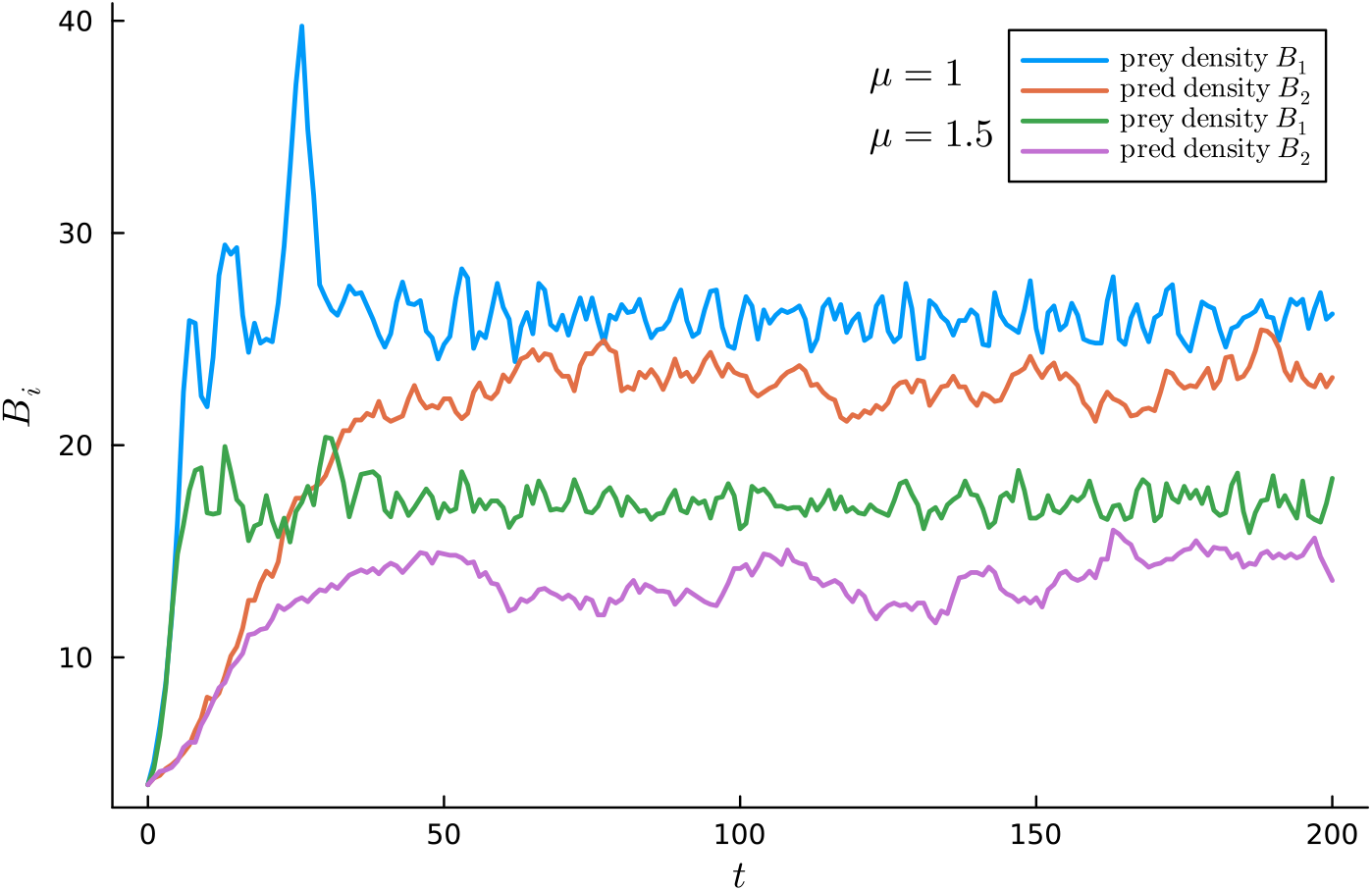
In the ABM, increasing the value of the metabolic rate *μ* decreases both predator and prey stationary density. The grid area is 4^2^, the initial density is 1 for prey and 10 for predators, the aggressiveness threshold is *ξ* = 500, the reproduction probabilities are *ω*_1_ = *ω*_2_ = 1/2, the catch probability is *δ* = 1/3, the resource growth-rate is *ρ* = 25, the predation efficiency is *η* = 10 and the metabolic rate is *μ* = 1 for blue and orange and *μ* = 1.5 for green and purple.

**Figure 14:**
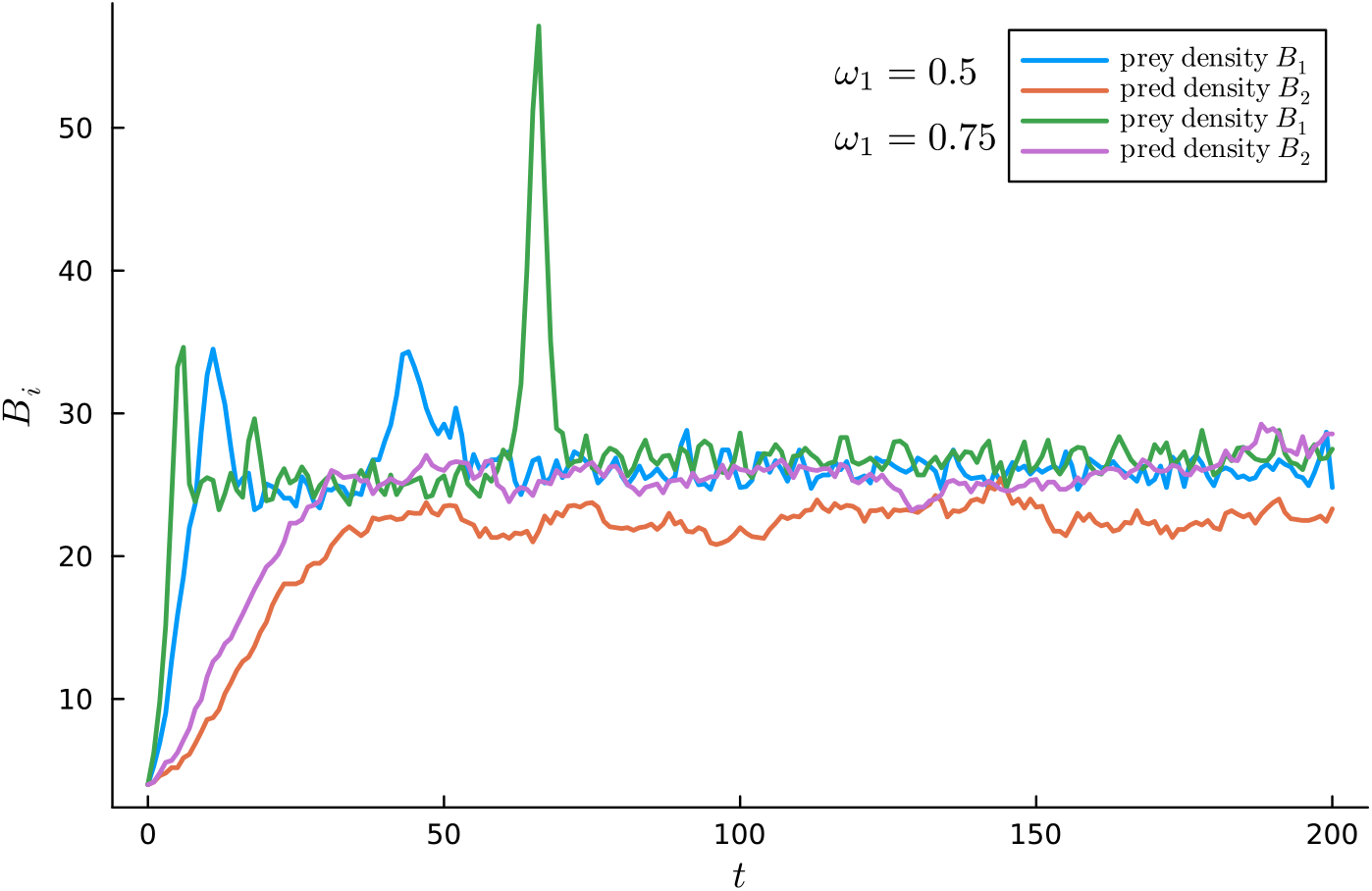
In the ABM, increasing the value of the prey reproduction probability *ω*_1_ increases predator stationary density and does not affect prey stationary density. The grid area is 4^2^, the initial density is 1 for prey and 10 for predators, the aggressiveness threshold is *ξ* = 500, the predator reproduction probability is *ω*_2_ = 0.5, the catch probability is *δ* = 1*/*3, the resource growth-rate is *ρ* = 25, the metabolic rate is *μ* = 1, the predation efficiency is *η* = 10 and the prey reproduction probability is *ω*_1_ = 0.5 for blue and orange and *ω*_1_ = 0.75 for green and purple.

The transition in which, under a certain resource regeneration rate, the predator population cannot be sustained, is a common phenomenology in predator-prey models. Consider, as an example, the system

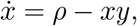

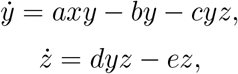

where *x, y* and *z* are respectively resources, prey and predator. The stationary solution for predators is

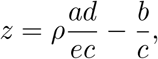

which means that no predator population can be sustained below *ρ* = *be/ad*.

## Footnotes

1 ^1^This exponent, incidentally similar to scaling laws found in complex systems outside of ecology [37], suggests that even a seemingly weak density-dependence may play a significant role in biomass patterns.

